# OmicLoupe: Facilitating biological discovery by interactive exploration of multiple omic datasets and statistical comparisons

**DOI:** 10.1101/2020.10.22.349944

**Authors:** Jakob Willforss, Valentina Siino, Fredrik Levander

## Abstract

Visual exploration of gene product behavior across multiple omic datasets can pinpoint technical limitations in data and reveal biological trends. The OmicLoupe software was developed to facilitate such exploration and provides more than 15 interactive cross-dataset visualizations for omic data. It expands visualizations to multiple datasets for quality control, statistical comparisons and overlap and correlation analyses, while allowing for rapid inspection and downloading of selected features. The usage of OmicLoupe is demonstrated in three diverse studies, including an analysis of SARS-CoV-2 infection across omic layers, based on previously published proteomics and transcriptomics studies. OmicLoupe is available at quantitativeproteomics.org/omicloupe

## Background

Omic analysis carries the potential to reveal new biological understanding and serve as a source of biomarkers. Still, omic data are challenging to work with, in part as they often contain considerable variation within and between experiments driven by both biological and technical factors, such as differing experimental conditions or sampling procedures. This variation needs to be considered to correctly interpreting the data. Furthermore, choices of algorithms and statistical procedures for processing the data cause additional differences in the final results [1,2]. The variation seen in the data can represent valuable biological trends, but can also be caused by nuisance factors, such as batch effects [3] or sample-to-sample technical variation. If the sources of trends in a dataset are understood, the dataset’s reliability can be assessed, and robust approaches of analysis and follow-up studies can be designed. Visualization is a critical tool for developing this understanding.

In comparative studies, one commonly overlooked aspect is the in-depth analysis of how individual features, such as transcripts or proteins, detected in one set of samples behave in other samples, datasets, or types of omics. Quality visualizations such as principal component analysis (PCA), and visualizations based on the outcome of statistical comparisons such as volcano plots and p-value histograms are often used to study trends within datasets. As an extension, several approaches to multiomics have been presented where the aim is to project down multiple sets of data to the same low dimensional space, such that they can be jointly visualized and inspected [4–6]. These provide useful overviews of multiomic datasets, but does not offer a detailed view of how individual features behave across multiple datasets in statistical comparisons.

Here, we propose an approach where single dataset visualization approaches are expanded to allow direct comparisons across datasets. Use cases are, for example, (1) Biomarker studies where an initial set of candidates is to be validated (2) Time-series experiment where the global expression is inspected, for instance, at different times after infection (3) Multiomics experiments where multiple types of data are produced for the same or similar biological systems and (4) Detailed studies of comparisons between methods or software approaches. To facilitate such analyses, we here introduce the interactive software OmicLoupe, which leverages additions to standard visualizations to allow for explorations of features and conditions across datasets beyond simple thresholds, giving insight which otherwise might be lost. The tool aims to be easy to use, directly interface with upstream software and to enable exploration and exporting parts of particular interest in the data. In the present work, we further demonstrate how OmicLoupe can be used to rapidly explore complex datasets in three different use cases

## Results

### Software implementation

To improve the accessibility and capability of analysis of complex datasets, we developed OmicLoupe. It is an interactive piece of software accessible through any web browser (quantitativeproteomics.org/omicloupe), which can either be accessed online or installed and launched locally as an R package (https://github.com/ComputationalProteomics/OmicLoupe). A Singularity container for execution without any prior dependency installation, and video tutorials are available to increase its accessibility. The code follows a modular design, promoting the extension of OmicLoupe with additional visualizations in the future.

OmicLoupe is built as a collection of modules, each performing a certain part of the analysis (Figure 1). It is built to fit into upstream workflows and can handle any combination of one or two expression datasets where the data are presented as tables with samples as columns and features (genes, proteins, transcripts or other measured features) per rows (illustrated in Supplementary Materials S1). Statistical visualizations require columns with p-value, false discovery rate (FDR) corrected p-values, fold change (difference of means between the two compared groups) and average feature level. These values are provided by up-stream software such as NormalyzerDE [7] or R packages such as Limma [8] for most types of omics or DESeq2 [9], for RNA-seq expression data. After loading the data in the web interface, the visualizations can be accessed immediately.

**Figure 1.**
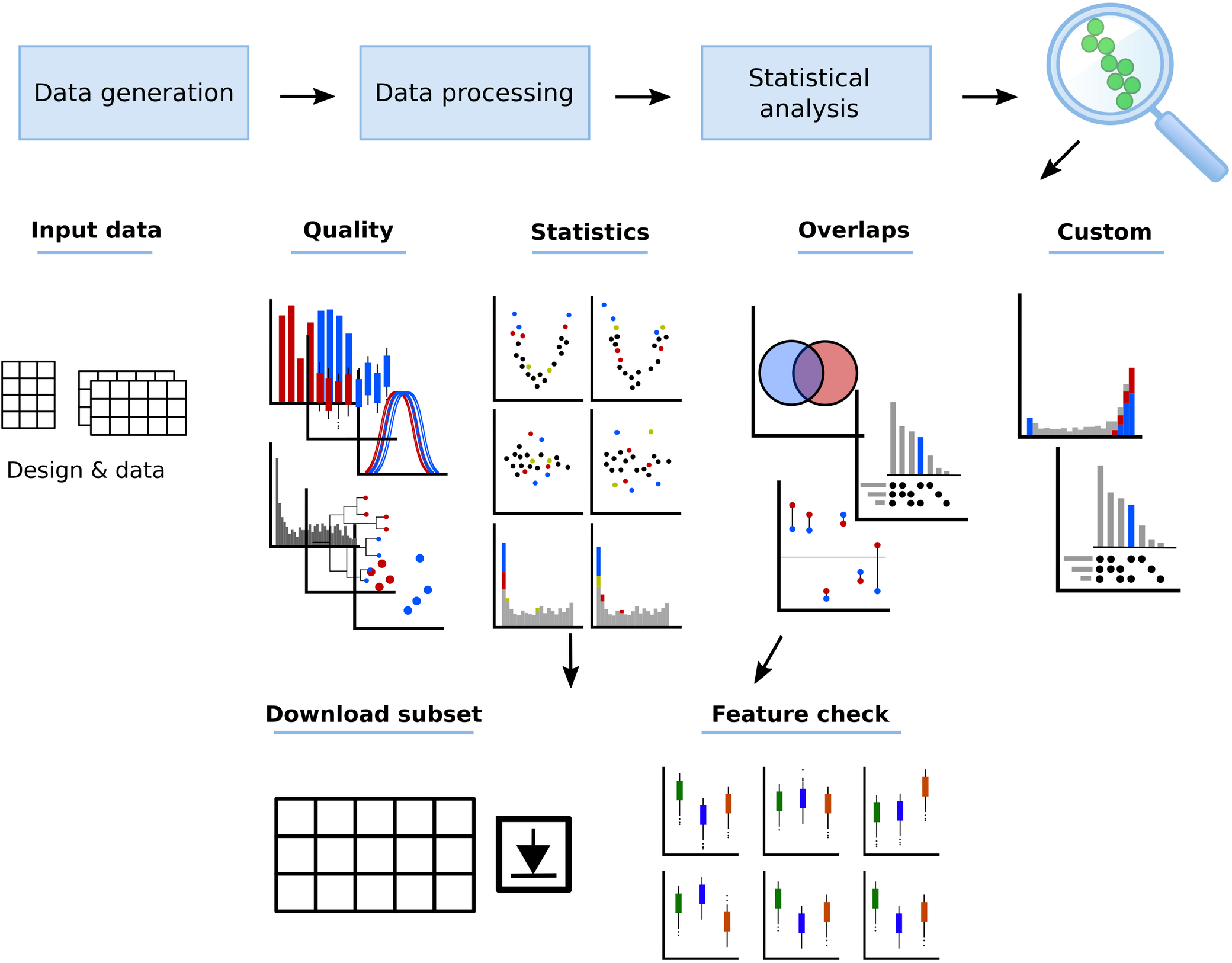
The OmicLoupe workflow. OmicLoupe is designed to easily interface with the upstream data generation process and to work on any expression data matrix. It provides the ability to explore up to two datasets, and provides comparative views between statistical contrasts performed either within one dataset or across multiple. It is organized in modules allowing rapidly shifting from a sample-wide view, to inspect individual statistical comparisons, overlaps between multiple comparisons, to understanding single features. (Adapted from schematics shown at http://quantitativeproteomics.org/omicloupe).

The general analysis workflow is shown in Figure 1. The workflow starts with the user assessing their data using the sample-wide quality visualizations, including boxplots, density plots, bar plots, dendrograms, histograms, and principal component plots. These visualizations commonly reveal outlier samples and the presence of systematic effects in the data. Further, for studies involving two datasets, OmicLoupe provides the side-by-side study of whether these effects are uniquely present in one or both of the datasets. These visualizations help the user to make decisions on how to best perform analysis such as outlier omissions, or decisions on what statistical comparisons to perform, and to judge the reliability of the data.

Next, the user can screen overlaps between statistical comparisons and sample conditions by inspecting whether features pass specific statistical cutoffs (p-value, FDR, optionally in combination with fold change) in one or several statistical comparisons (i.e. specific treatments or time points) or datasets. This overlap is illustrated by Venn diagrams for pairwise comparisons and UpSet plots [10] for a higher number of comparisons, with the UpSet visualization designed to efficiently compare a high number of overlaps. Further, for statistical comparisons, overlaps can be split by fold direction giving a better sense of whether overlaps indicate a shared abundance pattern. A novel visualization illustrates the fraction of features that change abundance in the same direction for low- and high-p-values, and the fold patterns of shared features are highlighted. These illustrations jointly provide a detailed view of similarities between contrasts. For both statistical and qualitative UpSet plots and for the Venn comparisons, subsets can be directly inspected and exported. Single features can be chosen for closer inspection in the Feature check module to evaluate in detail how their abundance values are distributed over any sample condition, shown either as raw data points or by using box- or violin plots.

Finally, to explore patterns across comparisons, the user can study pairwise comparisons and inspect how either significant or selected features in one dataset distribute across the volcano and MA scatter plots, as well as in p-value histograms. These visualizations can further be colored on any user-specified feature column and can be colored on respective loadings in the principal component plots, revealing groups of features strongly linked to specific principal components. Together, this yields an in-depth understanding of similarities and differences between contrasts and can display target features of interest. Similarly, for overlaps, single features can be selected and inspected in the Feature check module.

Beyond the previously mentioned, further specialized visualizations are provided. An analysis approach for identifying features uniquely present in certain conditions is provided as an UpSet plot, which can highlight features for which the abundance is below detection limit for certain samples. A correlation plot allows direct illustration of feature correlation patterns between data layers based on the same set of samples, for instance multiomics or alternative software processing of the same dataset. All the plots discussed above can be downloaded in PNG or vector format and can be customized, providing publication ready visualizations (Figures 2 to 7 in the present study are examples of visualizations generated using OmicLoupe). Furthermore, settings used at any point can be saved as a JSON file to ensure reproducibility (outlined in Supplementary S2 and available at https://doi.org/10.5281/zenodo.4110102 for the figures presented in this work).

**Figure 2:**
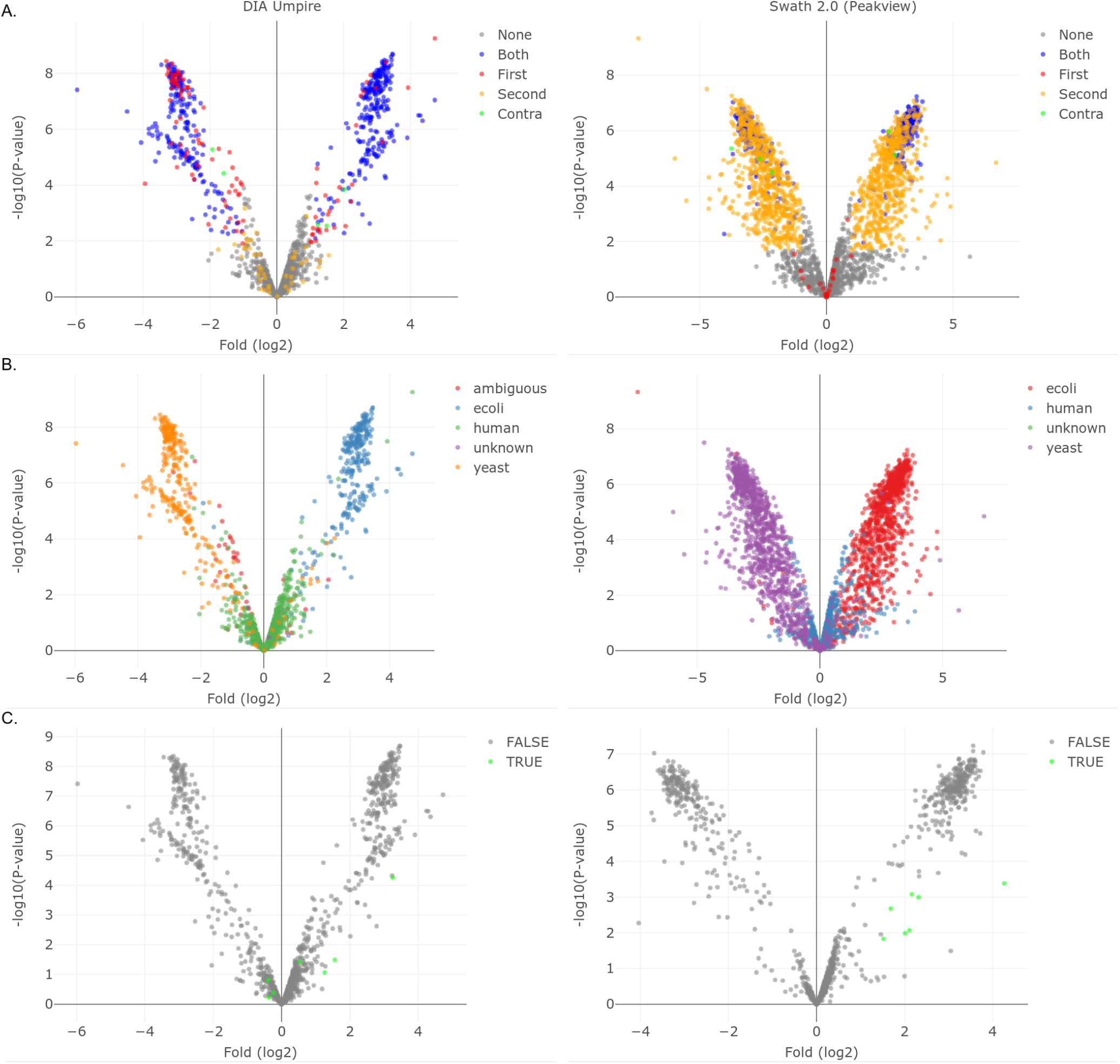
Effects of data processing using different software. Illustrations of data after processing with software: PeakView (pw), Skyline (sl), OpenSwath (os), DIA Umpire (du) and Spectronaut (sn). (A) Density distributions, available in OmicLoupe’s Quality panel, are colored on the data source (left) and spike-in level (right), respectively. (B) Features uniquely identified (not missing in all samples) after processing of different methods, with DIA Umpire highlighted. (B) UpSet plots of the features that were found as significantly differentially abundant (FDR < 0.05, log2 fold change > 1) when data had been processed using the different software. In C, features that are changing upwards or downwards in the comparison are displayed separately to visualize contradictory abundance changes due to differential processing. Eleven proteins that were deemed significantly changing, but with opposing direction of change after processing in PeakView and Spectronaut are highlighted in blue.

To summarize, OmicLoupe provides a tool to rapidly assess datasets for technical trends and for indepth studies of statistical comparisons and individual analytes.

### Case 1: Effects of data processing software on differential expression analysis outcome

To assess the utility and validity of the approaches introduced in OmicLoupe, we started by analyzing spike-in proteomic data that have previously been explored extensively in a comparison of data processing software for data-independent acquisition (DIA) LC-MS/MS data [1]. This dataset consists of *E. coli* and yeast proteins spiked at two different concentrations into a human proteome background. The two mixtures were analyzed in triplicates. In the original work, the data were processed using five different software, allowing for a comparison of their relative performance. The software used were PeakView, Skyline, OpenSwath, DIA Umpire, and Spectronaut, where only DIA Umpire was used without matching to a previously generated spectral library. This dataset was employed as an example with known ground truth where concentrations of proteins from different organisms were known, allowing assessment of how well the visualizations illustrate the known underlying trends. Further, it demonstrates how OmicLoupe can be used to assess the impact of different DIA software methods for processing the same set of samples.

Upon inspection in OmicLoupe, the quality control visualizations show that the choice of software impacts the resulting absolute values, as illustrated in the density plots shown in Figure 2A. Less obvious differences are seen between the spike-in levels, although upon inspection in a dendrogram, the difference between OpenSwath and PeakView appeared smaller than their respective spike-in levels (Supplementary S3). It can be noted that the intensity values were scaled to reach similar levels in subsequent analyses in the original study for this reason. Qualitative inspection was performed to identify proteins only detected by certain software processing methods. Here, the majority of proteins (2453) were detected by all five methods, and 1101 proteins were detected by all methods except DIA Umpire. Conversely, DIA Umpire identified 477 proteins that were not detected by any other method (highlighted in blue, Figure 2B), although PeakView identified a higher number of proteins uniquely (648, Figure 2B). Upon statistically comparing the abundances between the spike-in levels, a considerable number of proteins were also uniquely identified as differentially expressed by PeakView (488 spike-ins). Eleven proteins were found to be significant but with opposing direction of change when comparing PeakView and Spectronaut output (highlighted in blue, Figure 2C). Out of these, all eleven were yeast protein, correctly identified as downregulated by PeakView.

To further elucidate the underlying differences in the processed data from DIA Umpire and PeakView, a closer inspection of the statistical distributions was made in OmicLoupe. Out of the six statistical figures, the volcano plots are illustrated in Figure 3 (all six can be found in the Supplementary Material S4). Inspection of features passing the thresholds FDR < 0.05 and fold change (log2) > 1 (Figure 3A) shows a considerable number of common statistical features with the same abundance change directions (blue) but it is notable that a larger number of features are identified only after PeakView processing. These are distributed across all significance levels and folds, with no evident trends for a higher concentration of lower abundance values. A handful of features are found to be changing in opposite direction between the groups (green) and were compared to the expected regulation pattern. Here, it was found that all features with a positive log2 fold in both methods were true positives, while out of the two with negative fold-change, one was a false positive. This indicates how OmicLoupe could be used to qualitatively give indications of the reliability of features across comparisons by using fold change. Interestingly, sets of features found to be changing in one group are clustering around zero-fold change in the other method, indicating a different ability of the software to handle these features. Further inspection of the ground truth (Figure 3B) illustrates the respective types of spike-in. For PeakView, there seems to be, in particular for yeast, a considerable number of false negatives identified, while for DIA Umpire these are less common. To illustrate the joint use of the two methods, a subset of features identified after processing by both methods was inspected (Figure 3C). Here, a set of features, only identified as differentially abundant after PeakView processing (yellow in Figure 3A), was highlighted and their distribution after DIA Umpire processing inspected. One exception was significant in both cases, seen as the green point with lowest p-value (highest along the y-axis). Interestingly, the features represented a mix of true positives *(E. Coli)* and false positives (human). The true positives were found with greater fold change in DIA Umpire (two rightmost green points in Figure 3C, DIA Umpire panel). From these observations, we conclude that OmicLoupe allows for finegrained analysis of differences resulting from data processing using different software and allows careful inspection of specific data points across multiple datasets.

**Figure 3:**
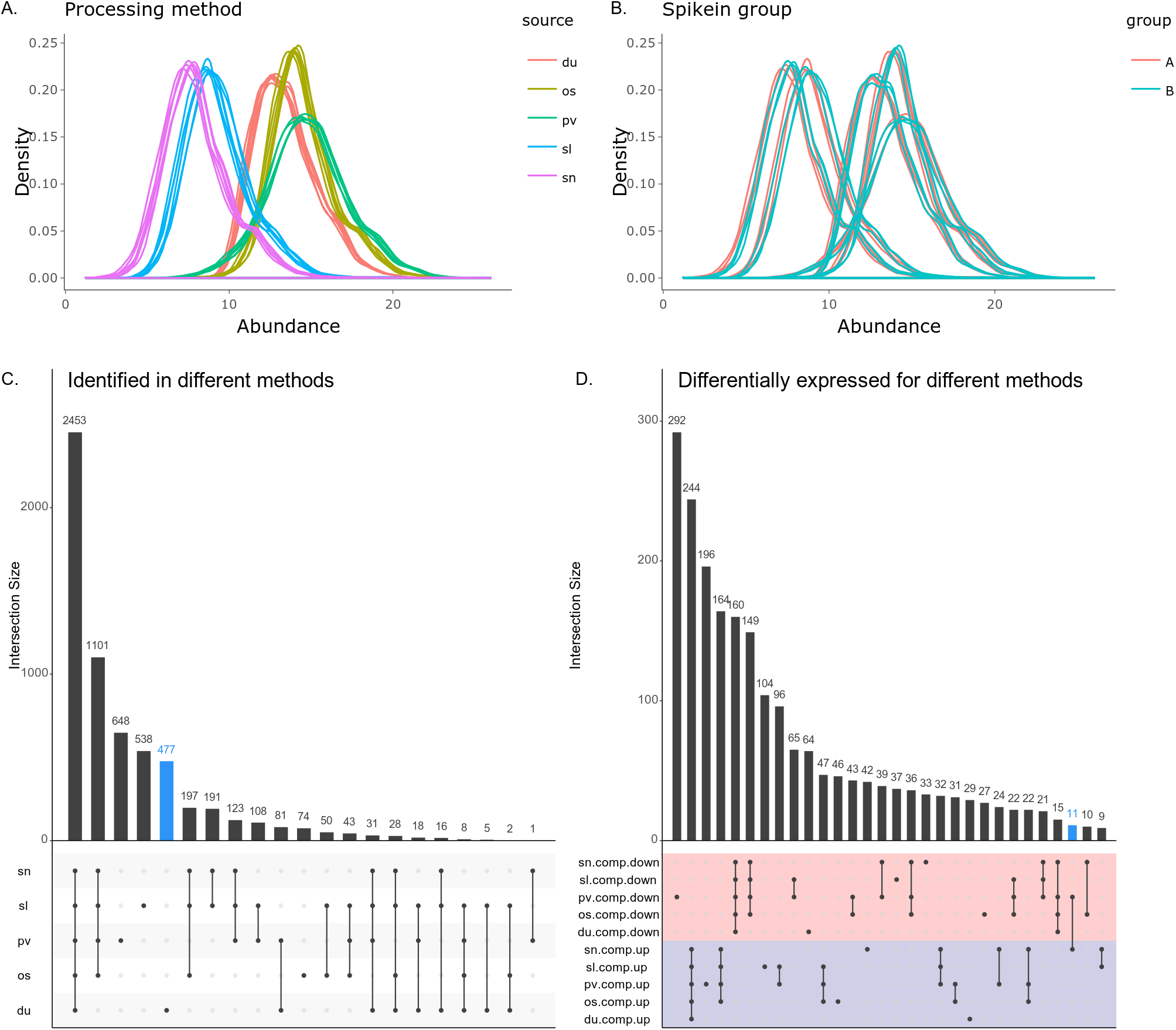
Comparison of contrast between spike-in levels in DIA Umpire and PeakView. These panels show part of the statistical interface in OmicLoupe. Panel A) shows features passing the significance threshold FDR < 0.05 and log2 fold > 0.5 in individual datasets, and in both. Green points (“contra” in the legend), are passing the significance threshold in both datasets, but with reversed log2 fold direction. Panel B) shows coloring based on the spike-in source. Panel C) show the outcome of interactively highlighting a set of five features only significant in PeakView and one significant in both. This reveals their distribution in DIA Umpire, showing that the features upregulated in both are true positives, while one of the two found in lower abundance in DIA Umpire is a false hit.

### Case 2: Analysis of matched proteome and transcriptome data

Multiomics studies of the same biological samples are becoming increasingly more frequent, but how to integrate the data types and finding important features remains challenging. We thus investigated how OmicLoupe can be used for direct comparisons of different data types taken from the same set of samples, to reveal features only detected in certain conditions, and common patterns of observed abundance level changes. For this purpose, a comprehensive multiomics dataset from endometrial cancer samples was downloaded [11]. Multiple types of data, including proteomics and RNA-seq, were acquired for the samples in the original study, and the features had been mapped to common gene identifiers. The samples are classified in different histological types, including Copy Number Variation (CNV) high, which includes serous samples and cancers penetrating at least 50% of the endometrial wall and CNV low consisting of samples penetrating less than 50% of the endometrial wall. Here, the statistical analysis focused on the comparison between these groups.

A first view using the PCA module revealed a primary separation between most normal samples and tumor samples (Figure 4A). This separation was similar in both proteomics and RNA-seq datasets, with few noticeable differences. Further, a partial separation between CNV high and CNV low can be seen along with the second principal component. PCA analysis without the control samples group was also performed using the function available in OmicLoupe (Supplementary S5). To study the similarity of the statistical comparisons across the two data types, features with positive abundance change and with low p-values were highlighted in the RNA-seq contrast (by dragging directly in the figure) between CNV high and CNV low to see how these distribute in the corresponding contrast in the proteomics dataset (Figure 4B). The majority of features were also upregulated in the proteomics data with three exceptions, namely FOLR3, STEAP1B and TBL1D31. Genes of interest, including TP53, which was discussed in the original study, were inspected using the Feature check (Figure 4C), which showed a reversed pattern in transcriptomics and proteomics. Finally, the correlation between transcriptomics and proteomics was studied. Pearson and Spearman correlations are illustrated in Figure 4D and showed similar median values (0.51 Pearson) to those presented in the original study, with a small number of inversely correlated features.

**Figure 4:**
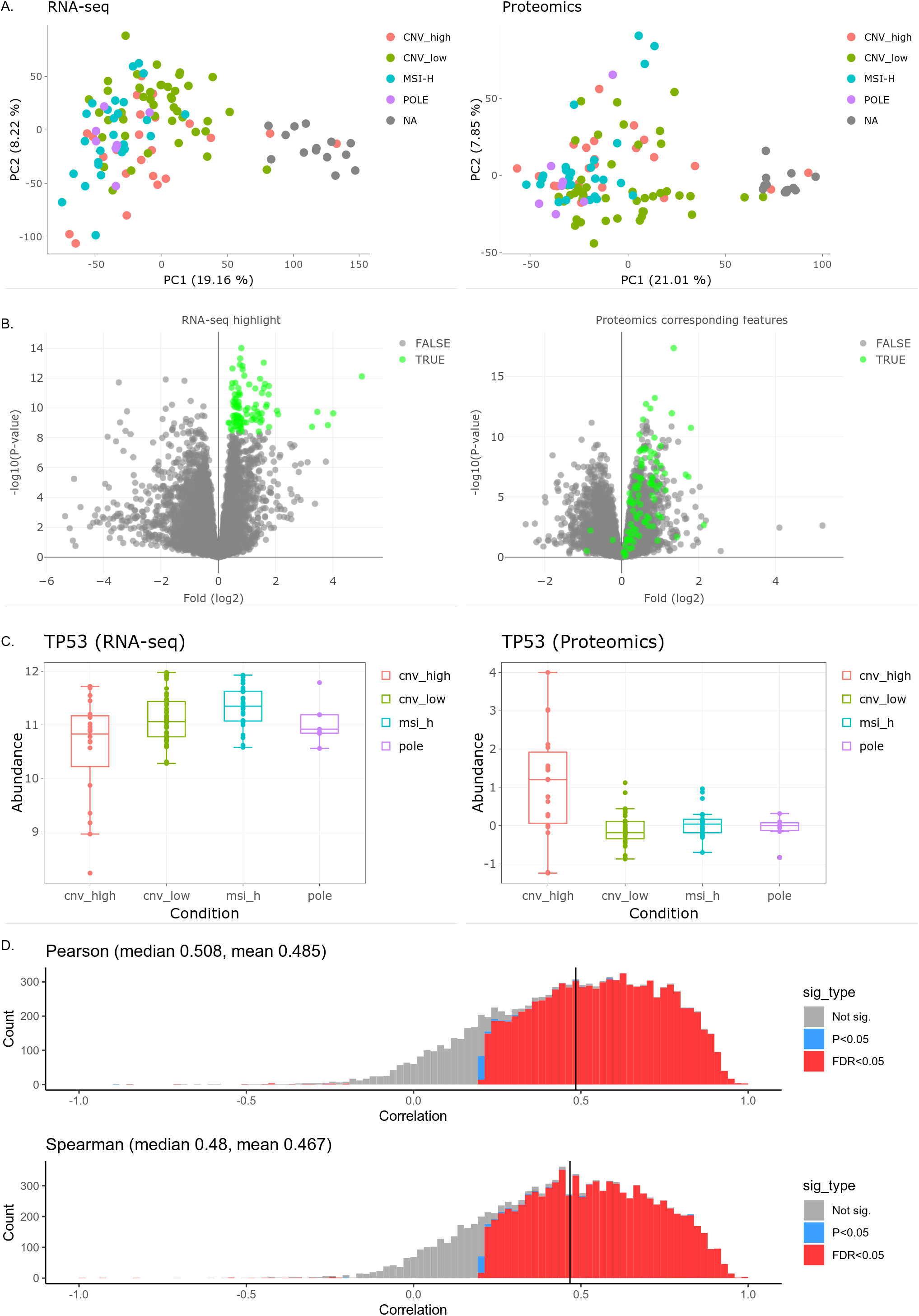
Illustrations of multiomics data investigated in Case 2. A) Principal component illustration present in the quality module, comparing proteomics and transcriptomics. As can be seen, the major trends are similar between the two data types. B) Distribution of how high-significance features upregulated in RNA-seq distribute in the proteomics dataset. C) Boxplots of TP53, identified in the dataset across the four studied sample classifications using the feature check module. D) Correlation distributions between the RNA-seq and proteomics features using the correlation module.

This demonstrates how OmicLoupe can be used to inspect similarities and differences between layers of omic data generated from the same set of samples, providing an improved understanding of both the general expression profiles and individual gene products.

### Case 3: Cross dataset comparisons of proteome and transcriptome data from two published SARS-CoV-2 studies

When studying proteins, for instance involved in certain diseases, validation of key proteins in multiple experimental setups can provide valuable biological information. However, it can be demanding to handle omic data from different studies. We aimed to assess whether OmicLoupe could facilitate this process and aid in finding shared key features. Herein, we examined SARS-CoV-2 infection of two model systems from different studies with time series data to find overlapping features. The first study employed human colon epithelial carcinoma cell line (Caco-2) infected by the virus at different time points (2, 6, 10 and 24 hours), with corresponding control samples, and with both proteomics and translatomics read-out (Bojkova *et al*., 2020). In the second study [12], the transcriptomic analysis had been performed on human intestinal organoids, representing the human gut, infected with the virus at time points 0, 24 and 60 hours and in two different media: differentiation and expansion media. The datasets were first overviewed individually using OmicLoupe and then analyzed jointly to find common patterns.

For the proteomics study, the initial quality control revealed two aspects in the data influencing the subsequent analysis. Quality control visualizations using PCA and dendrograms revealed clustering of samples according to a sample name-based categorization, thought to be the plating numbers of the cell lines (Figure 5A). To compensate for this effect, this number was included as a covariate in the statistical tests, and the impact of including it was investigated in OmicLoupe. The inclusion of this covariate led to a considerably higher number of features detected as significantly different at the thresholds employed (FDR < 0.05 and log2 fold change > 0.3), as exemplified for the comparison of infected samples at 6 hours versus infected samples at 2 hours (Figure 5B). Next, OmicLoupe was used to study control and infected samples independently (as illustrated in Supplementary S6). Here, a clear pattern was seen in the infected samples, with the 24 hours infected samples separating out along PC1, while the 10 hours samples showed a weaker separation. In the control samples, the trend was less clear, and the control 6 hours samples appeared as weak outliers. In order to study the potential impact of these group comparisons, control and infected samples at 6 and 2 hours were compared, as depicted in Figure 5C. A strong effect of decreasing abundance is seen in control 6 hours, while in infected 6 hours the trend is smaller, with known viral proteins being clearly upregulated. This unexpected distribution of the 6 hours control samples led us to focus on comparisons between infected samples, and the 24 hours infected versus control comparison.

**Figure 5:**
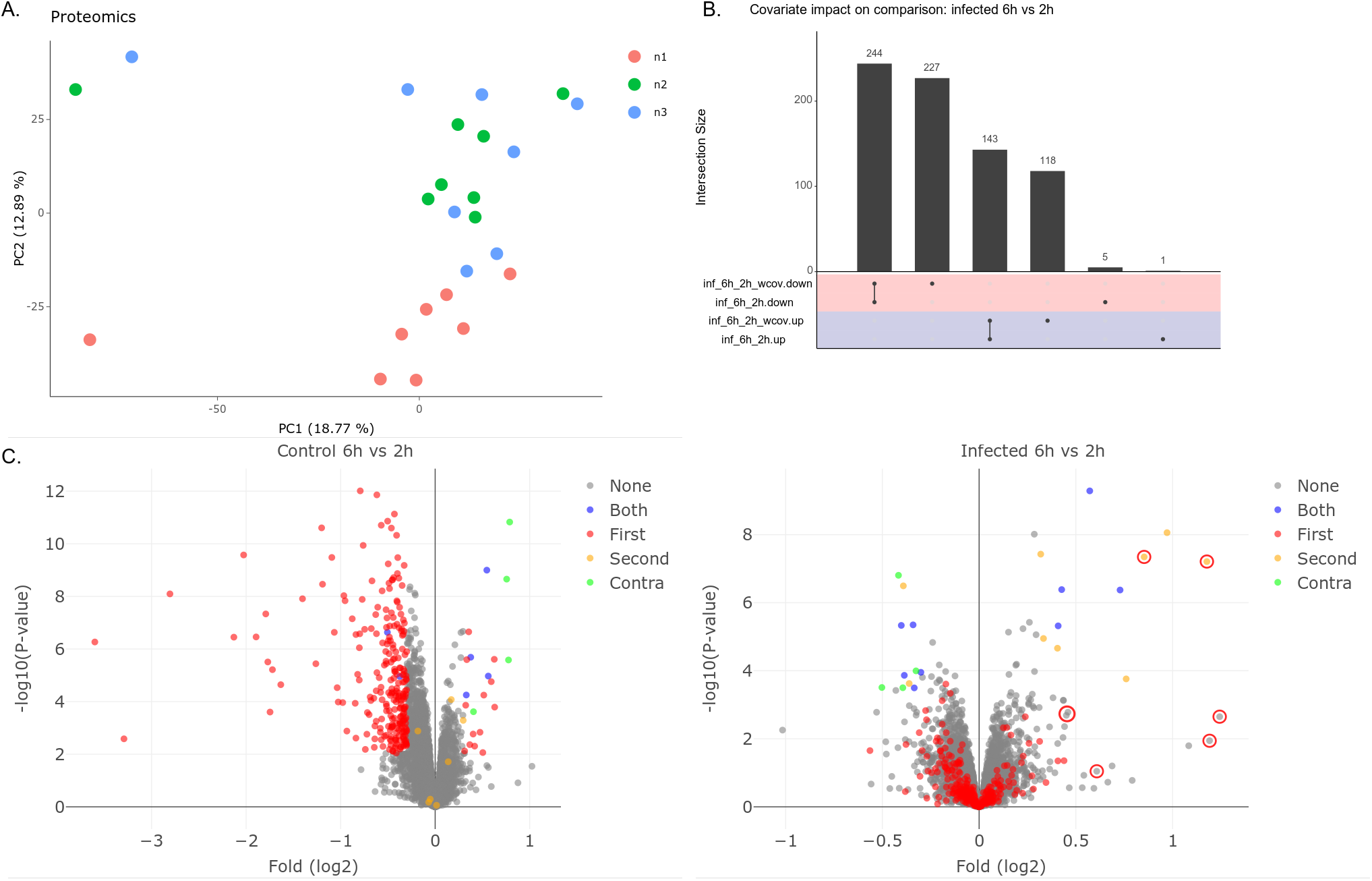
Quality inspection of the SARS-CoV-2 proteomics dataset. A) Inspection revealed a separation of samples along the second principal component likely related to a plating effect. This was compensated for in subsequent statistical tests by including it as a covariate. B) The impact of performing differential expression analysis without and with including the putative plating number as a covariate. The inclusion of the covariate yielded 345 new statistical features while losing six as compared to not including the covariate. C) Comparison of control samples 6h and 2h shows many features with a decrease in abundance, indicating that the mock treatment might influence the data. Comparison between infected samples at 6h and 2h show more limited differences, with seven detected viral proteins among those with increased abundance at 6 hours indicated in red circles (log2 fold change > 0.3) out of which two passed the FDR threshold.

To study the viral distribution between the infected conditions we highlighted proteins with known annotation related to either virus or virus receptors (According to UniProt https://covid-19.uniprot.org, downloaded 2th of July 2020) in comparisons of 6, 10 and 24 hours infected samples compared to 2 hours infected samples. Figure 6A clearly shows how the viral proteins are increasing in abundance in infected cells at 10 hours and even more at 24 hours. This first overview serves as quality control and is in line with what was shown in the original study. For the transcriptomic dataset, using the same cutoff parameters used for the proteomic study, a first evaluation shows that at 24 hours after infection in the differentiation media, 654 transcripts are differentially expressed, while in the expansion media after 24 hours of infection, 438 transcripts are differentially expressed.

**Figure 6:**
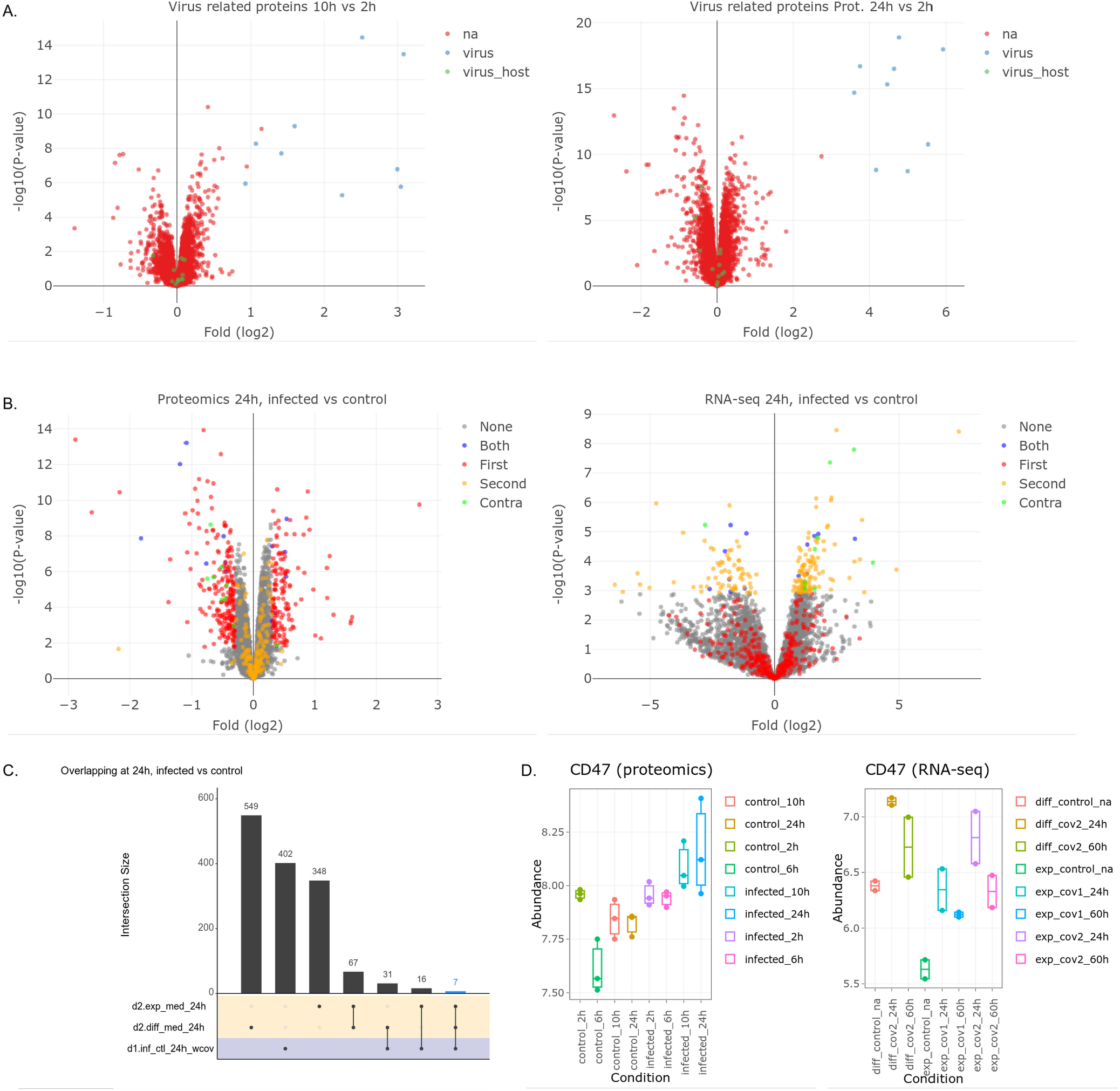
Inspection of trends in SARS-CoV-2 proteomics and transcriptomics datasets. Statistical analysis performed using the following settings: FDR < 0.05 and log2 fold change > 0.3. For both datasets, data from 24 hours after viral infection are used A) Inspection of infected samples, 10 hours and 24 hours compared to 2 hours, colored by proteins known as virus proteins and virus receptors revealed a clear upregulation among virus proteins. B) Direct comparison of infected and control at 24 hours after infection in proteomics and comparison between the proteomic dataset and the transcriptomic dataset (expansion media). The coloring is based on in which dataset the gene products pass the significance threshold. Green points (“contra” in the legend), are passing the significance threshold in both datasets, but with reversed log2 fold direction. This comparison revealed a set of shared proteins, both changing abundance in the same or opposite direction. C) Illustration of the shared significant genes between proteomic and transcriptomic (differential and expansion media) datasets. D) CD47 distribution at different time points in the proteomic study and in the transcriptomic data.

Furthermore, to study the potential of OmicLoupe, the results from the proteomic study were compared to the transcriptomic dataset. Here, the two datasets can easily be uploaded and compared based on their time points. To make a similar comparison in the two datasets, we decided to compare the proteomic and the transcriptomic data at 24 hours, despite one outlier sample being identified in the transcriptomics data at this time point (the PCA plot for the transcriptomics data illustrating this outlier is shown in Supplementary S7). The distribution of the significant genes in the proteomic study in comparison with the transcriptomic study (expansion media), is depicted in the volcano plot in Figure 6B. Of particular interest are the significant genes that are shared between the datasets at 24 hours after infection. At the set threshold (FDR < 0.05 and log2 fold change > 0.3), 38 differentially expressed genes are shared between the proteomic and the transcriptomic data after 24 hours of infection in the differentiation media. For the extension media, 23 genes are significantly shared between the proteomic and transcriptomic datasets after 24 hours of viral infection. The overlap between the IDs in both datasets is displayed in the UpSet plot in Figure 6C. Interestingly, 7 genes were overlapping between the proteomic dataset and the transcriptomic study (differential and expansion media). As an example, one of those shared genes, CD47, is depicted in figure 6D. CD47 is a leukocyte surface antigene, which has been shown to be upregulated after a viral infection, including SARS-CoV-2 infection, as a host response to the infection [13]. These overlapping groups were further analyzed in STRING [14] to investigate the relevant pathways connected to these significant genes identified. Of biological interest is that one of the main regulated Reactome [15] pathways is neutrophil degranulation, which in many studies has been reported as a key biological process during the SARS-CoV-2 infection [16–18].

In summary, in this third case study, OmicLoupe was used to perform a parallel analysis of two datasets from different types of omics (proteomics and transcriptomics) to investigate the response to infection over time. Both these datasets were obtained from published studies. By straightforward visualizations, we demonstrate the feasibility of using this tool to easily identify significantly changing gene products, common to both datasets, which can be used for further analysis, such as GO enrichment and pathway analysis.

## Discussion

Visualization is an important tool to fully explore omics datasets and to highlight features that can be difficult to assess with numbers alone. Consequently, there are many new software solutions for omic data visualization presented over the past few years. These include a range of user-friendly stand-alone software for omics visualization such as Perseus [19] for proteomics, or shiny-based software such as ShinyOmics [20], which provides a flexible quality-oriented interface to omic data, and WIlsON [21] providing high-quality interactive figures based on an open file format but only limited abilities to compare features. Intervene is a software focusing on comparisons [22], aiming to provide various types of overlap information, but only based on fold change information and not allowing for feature-by-feature examination. Furthermore, software dedicated to incorporating multiple layers of omics such as MixOmics [23] has extensive multiomics integration capabilities, but does so on a sample-wide scale rather than focusing on the behavior of single features. Despite these notable examples and several more new visualization software packages now being available, we have failed to find software performing several of the functionalities for multiple comparisons across datasets provided by OmicLoupe. Key features in OmicLoupe like the side-by-side data distribution comparison volcano and MA plots, with labeling of key features across the comparisons and of datasets, as well as the ability to rapidly switch to individual feature views across samples, enable a deeper understanding of the individual features in the data.

The implementation of UpSet plots with optional splitting based on changes in abundance direction, can rapidly help in determining reproducibility across datasets. While standard statistical comparison, using strict thresholds in many cases is the default option, underlying trends can be found in plots such as UpSet with less strict thresholds, when the data are lacking power.

Here, we explored three diverse datasets to highlight different aspects of OmicLoupe’s functionality. By comparing the impact of different proteomics software processing methods, we could study in detail differences in outcome between the methods, and identify specific features handled correctly only by one or some of the methods. Next, multiomics exploration with both transcriptomic and proteomic data obtained from the same samples gave the opportunity to explore features across omic layers. Here, we identified an overall similarity of trends across the omic data layers, and rapidly illustrated the correlations of transcripts and proteins. Further, we visualized key features in detail, including TP53, a key protein discussed in the original study, and detected differences at transcript and protein level. This demonstrates how OmicLoupe can confirm and provide extended knowledge for existing data. Finally, we used two separate SARS-CoV-2 studies to profile intestine cells during infection. OmicLoupe was used to identify and navigate technical limitations, including a batch effect, and a seeming lower reliability of one set of control samples. The overall regulation patterns were relatively different, as expected due to the different types of samples, but still subsets of features with joint abundance changes were identified. These were downloaded and enriched, revealing biological trends in line with what had been observed in prior studies.

The cases presented in this manuscript are common examples of challenges encountered when analyzing omics data. Beyond these, the OmicLoupe software has the potential to be used in a wide range of scenarios, to better understand both single- and multiple-omics datasets. To this end, we believe usability is of critical importance for this kind of software, and OmicLoupe has a straightforward interface, with user help text complemented with video tutorials at the website. Having these at hand may mean the difference in how extensively the data could be explored, and thus how well they can be understood. We thus encourage users to test the software, provide feedback about its functionality and to comment on possible useful new extensions.

## Conclusions

Here, we have presented OmicLoupe, which both introduces novel approaches for comparative visualization cross dataset and presents these in an interactive easy-to-use software. We have demonstrated its utility on three diverse datasets, starting with a technical dataset to demonstrate how OmicLoupe can be used for comparing processing methods and how the cross-comparison fold can provide important information. Secondly, we explored a multi-omic cancer dataset illustrating how same-sample cross-omics can be readily illustrated. Finally, to demonstrate its versatility, we reanalyzed two recently published SARS-CoV-2 datasets, performed comparative explorations of these datasets and rapidly identified proteins and RNA transcripts showing the same abundance change trends across both studies. Based on these results and usage on other datasets, we propose that OmicLoupe can be a versatile tool in many expression omics-based analyses, both for novice and expert users. We provide it for usage by the community, as an R-package and as an online server.

## Methods

### Development

OmicLoupe is implemented using R (v3.6.3) and RShiny (v1.4.0.2), using packages providing interactive visualizations: Plotly (v4.9.2.1), DT (v0.13) and packages for data visualization: ggplot2 (v3.3.0), GGally (v1.5.0), UpSetR (v1.4.0, [10]) and dplyr (v0.8.5) for data processing. The code is developed in modules to facilitate reusability. Further, a Singularity container [24] was prepared allowing immediate local execution without being required to install the R package dependencies.

### Dataset analysis

OmicLoupe was evaluated using three datasets covering different use cases. An R notebook containing the code used for preprocessing the datasets together with an HTML-document with the code output is outlined in the Supplementary S8 and accessible on the DOI: https://doi.org/10.5281/zenodo.4110102.

### Case 1: Technical spike-in dataset

A technical dataset was employed where proteins from human, *E. coli* and yeast had been spiked in at controlled concentrations [1] and subsequently analyzed using five different DIA methods. The data was downloaded from ProteomeXchange [25] at the ID PXD002952, selecting the data generated on the TripleTOF 6600 instrument with 32 fixed-size windows for all five methods. The HYE110 dataset was used, with a spike-in difference of log2 fold 3. The raw data matrices were preprocessed both into five separate data matrices, and into a single merged matrix consisting of all joint protein entries. They were subsequently log2-transformed and rolled up to protein level using an R-reimplementation (github.com/ComputationalProteomics/ProteinRollup) of the DanteR RRollup [26], using default settings and excluding proteins supported by a single peptide. Statistical contrasts between the two concentration levels were subsequently calculated using Limma [8] as provided by NormalyzerDE [7], and resulting p-values adjusted for multiple hypothesis testing using the Benjamini-Hochberg procedure [27].

### Case 2: Multiomics dataset

A multiomics dataset from a study investigating 95 prospectively collected endometrial carcinoma tumors [11] divided into four histological groups, including copy number (CNV)-high (serous cancer - a rare aggressive variant, and cancers with more than 50% penetrance of the endometrial wall) and CNV-low (less than 50% penetrance of the endometrial wall). The data matrices and meta information were downloaded from the supplementary information of the original study. Samples omitted from the original study were similarly omitted, as specified in the matrix obtained from the original supplementary. Further, upon inspection with OmicLoupe the set of normal samples was identified as a strong outgroup and omitted to avoid influence in the statistical procedure. The original dataset contains multiple layers of omic data, out of which the following two were used in the present study: proteomics and mRNA levels. For the proteomics matrix, statistical contrasts were calculated using Limma [8] in NormalyzerDE and Benjamini-Hochberg [27] corrected FDR values were calculated. For the transcriptomics matrices the data was provided as RSEM estimated counts. It was first transformed with Voom with quality weights [28] and subsequently processed using Limma [8]. Statistical contrasts were for both data types calculated between samples classified as CNV-high and CNV-low.

### Case 3: SARS-CoV-2 datasets

The first dataset analyzed in this case study is a recently published SARS-CoV-2 proteomic dataset [29], where human colon epithelial carcinoma cell line (Caco-2) was infected by SARS-CoV-2 and proteomic analyses were performed on samples at four time points (2, 6, 10, 24 hours after infection), both for infected samples, and samples treated with a mock infection. The proteomic data and metadata were generously provided by the authors in the supplementary materials of the study Zero-values were replaced with NA, and the protein abundance values were log2 transformed. Statistical contrasts were calculated using Limma [8] in NormalyzerDE [7] and resulting p-values FDR-corrected using the Benjamini-Hochberg procedure [27]. Initially, statistical comparisons were made between infected and control samples at each of the four time points (2, 6, 10 and 24 hours after infection). After initial explorations in OmicLoupe a batch effect was identified, which was subsequently included as a covariate in the statistical test, as described in the results section.

The second dataset was from human intestinal organoids infected with SARS-CoV-2 in both differentiation and expansion media and analyzed at two time points after infection (24 and 60 hours) using transcriptomics [12]. The data was retrieved from the NCBI Gene Expression Omnibus (GEO) database, from the accession number GSE149312. The data were TMM normalized, and Voom transformed [28] with quality weights. Subsequently, statistical calculations were carried out using Limma [8], comparing infected samples at 24 and 60 hours after infection to the uninfected reference. P-values were FDR-corrected using the Benjamini-Hochberg procedure [27].

## Declarations

### Ethics approval and consent to participate

Not applicable

### Consent for publication

Not applicable

### Availability of data and materials

- On its main page (http://quantitativeproteomics.org/omicloupe), a public server running OmicLoupe can be accessed, as well as links to video tutorials.
- The R package and its source code is available at github.com/ComputationalProteomics/OmicLoupe with DOI: https://doi.org/10.5281/zenodo.4113746
- A singularity container is available at singularity-hub.org/collections/4795 which allows execution with no need to install dependencies.
- The data used for case study 1 is accessible through the ProteomeXchange database under ID PXD002952 (http://proteomecentral.proteomexchange.org/cgi/GetDataset?ID=PXD002952)
- The data used for case study 2 is accessible from the study by Dou *et al* [11] (https://doi.org/10.1016/j.cell.2020.01.026) and is based on Supplementary Tables 1 and 2.
- The proteomics data used for case study 3 is accessible from the study by Klann *et al* [29] (https://doi.org/10.1038/s41586-020-2332-7) and is based on Supplementary Tables 1 and 2.
- The transcriptomics data used for case study 3 is accessible through the Gene Expression Omnibus database under ID GSE149312 (https://www.ncbi.nlm.nih.gov/geo/query/acc.cgi?acc=GSE149312)
- The settings used in OmicLoupe to produce all visualizations in the manuscript and supplementary materials are available at https://doi.org/10.5281/zenodo.4110102 and outlined in Supplementary S2.
- The R code for processing these datasets into a format compatible with OmicLoupe is available at https://doi.org/10.5281/zenodo.4110102 and outlined in Supplementary S8. These output matrices are available from the corresponding author on reasonable request.

### Authors’ contributions

The study was conceived by JW and FL. The software development was carried out by JW with input about functionality from VS and FL. Data analysis was performed by JW and VS. The manuscript was drafted by JW and was expanded, edited and approved by all authors.

## Acknowledgements

We want to thank Victor Lindh whose master’s thesis project allowed us to explore ideas which later were further developed here. We also thank OmicLoupe users who have tested and provided valuable feedback. Finally, we would like to thank the authors behind the four datasets used in the three case studies who generously provided their data for further use in an accessible way.

