## Supplementary for "OmicLoupe: Facilitating biological discovery by interactive exploration of multiple omic datasets and statistical comparisons"

#### Table of content


- Supplementary materials S1: Format for input tables
- Supplementary materials S2: OmicLoupe settings retrieved in JSON format for figures presented in the manuscript and in the supplementary materials
- Supplementary materials S3: Case 1, dendrogram illustration of impact from software processing methods
- Supplementary materials S4: Case 1, full statistical panel illustration of DIA Umpire compared to PeakView
- Supplementary materials S5: Case 2, PCA illustrations for subsets excluding non-cancerous data
- Supplementary materials S6: Case 3, proteomics PCA illustration of control- and infected samples
- Supplementary materials S7: Case 3, PCA illustration of transcriptomics samples
- Supplementary materials S8: Outline of analysis code as an R-markdown document together with output in HTML format

#### Supplementary materials S1: Format for input tables

The design matrix should contain one column with sample names that exactly match the sample columns as present in the data matrix. Beyond this column, it can contain multiple columns with biological and technical conditions. These are used for visualizations in OmicLoupe.

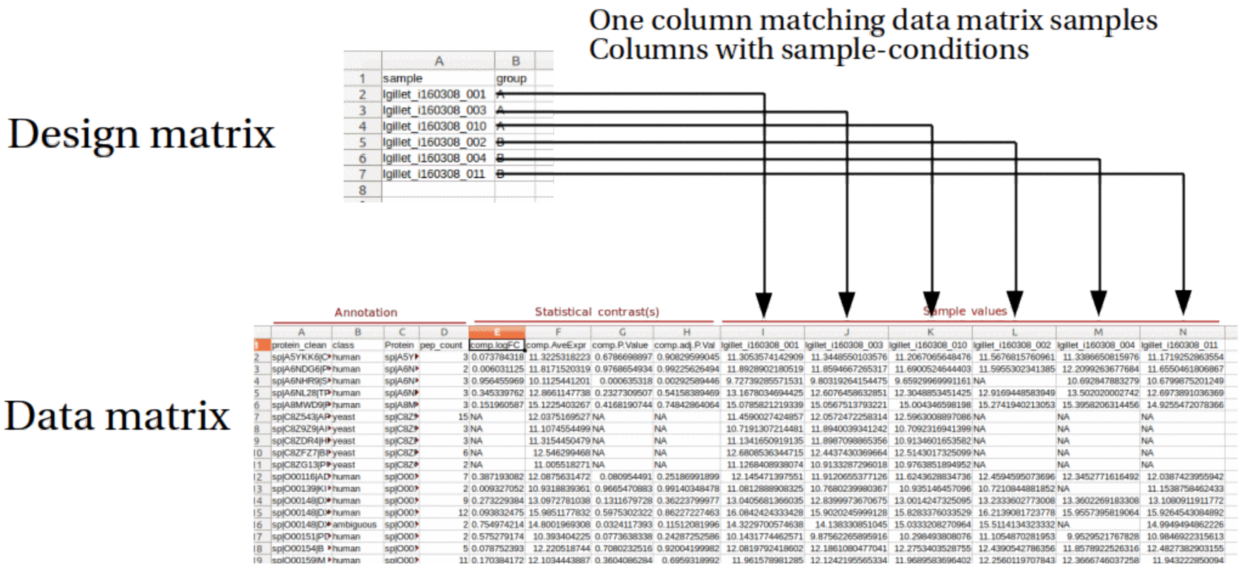


The data matrix can contain data columns, annotation columns and statistical columns. To include statistical contrasts, four columns per contrast needs to be included in the data matrix. These values include: The average expression for the feature (AveExpr), the log-scale difference between the compared conditions measurements (logFC) and the resulting p-values (P.Value) and adjusted p-values (adj.P.Val).

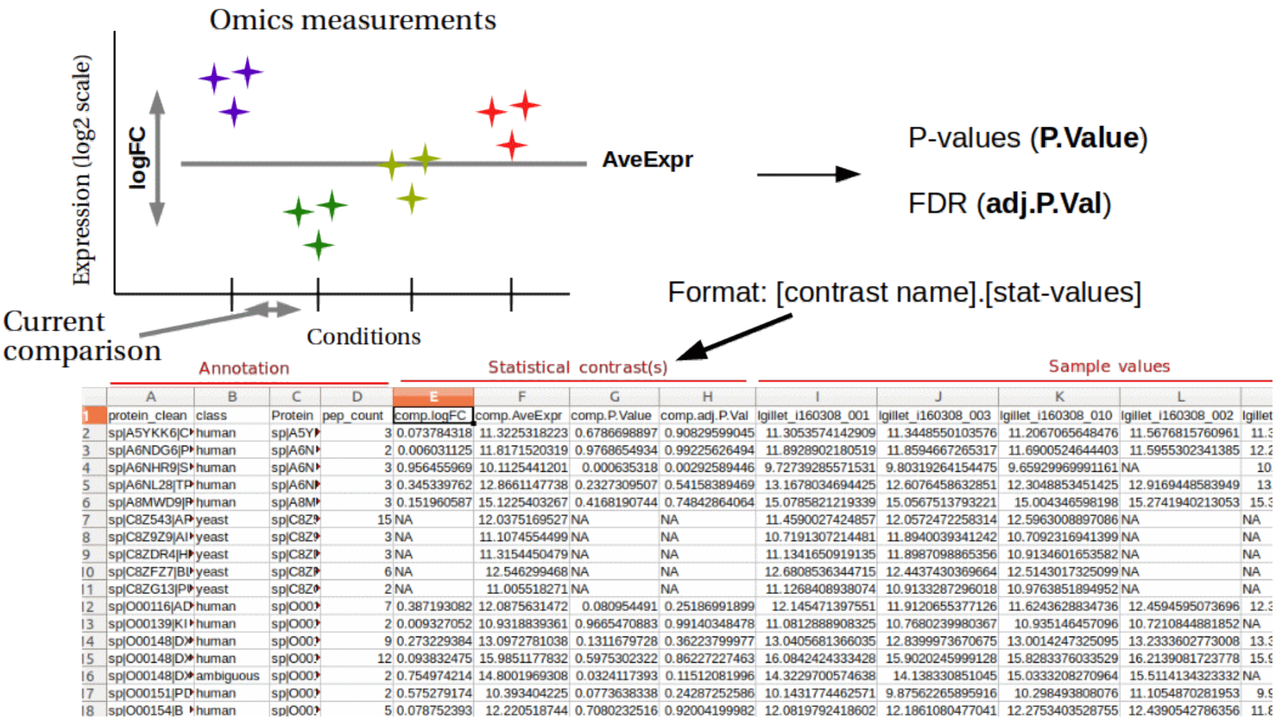

If using multiple data tables (can be different omics, as shown below, or from the same type of omics), one column needs to contain common IDs for the two cases (for instance, gene IDs).

Proteomics

| gene_symbol | accession | species_names | featureAvg | infected_2h-control_2h_PValue | infected_2h-control_2h_log2FoldChange | infected_2h-control_2h_AdjPVal |
|---|---|---|---|---|---|---|
| AFP | P02771 | Homo sapiens OX=9606 | 11.031331403655 | 0.30969902158687 | -0.307359232716664 | 0.99980939482126 |
| FABP1 | P07148 | Homo sapiens OX=9606 | 10.954591559394 | 0.587573151279121 | -0.209476148787799 | 0.99980939482126 |
| VIL1 | P09327 | Homo sapiens OX=9606 | 10.698272236802 | 0.416645112976052 | -0.191097088944636 | 0.99980939482126 |
| KRT18 | P05783 | Homo sapiens OX=9606 | 11.2095110506113 | 0.633331356317085 | -0.132690067072895 | 0.99980939482126 |
| | Q9P2E9 | Homo sapiens OX=9606 | 10.4230791312684 | 0.459488934884819 | -0.167897844503694 | 0.99980939482126 |
| ANXA4 | P09525 | Homo sapiens OX=9606 | 10.6499881385456 | 0.82781011607852 | -0.050223942845536 | 0.99980939482126 |
| KRT8 | P05787 | Homo sapiens OX=9606 | 10.619917799405 | 0.844472953537969 | -0.060373667476336 | 0.99980939482126 |
| MUC13 | Q9H3R2 | Homo sapiens OX=9606 | 9.37852849994093 | 0.831812491341271 | 0.078448770880204 | 0.99980939482126 |
| CDH17 | Q12864 | Homo sapiens OX=9606 | 9.86333520601928 | 0.518411935369265 | -0.142734075113003 | 0.99980939482126 |
| LGALS3 | P17931 | Homo sapiens OX=9606 | 10.2031600815392 | 0.489722341053065 | -0.135236521146233 | 0.99980939482126 |
| SERPINA1 | P01009 | Homo sapiens OX=9606 | 10.0902246911767 | 0.55887555105166 | -0.136333152852051 | 0.99980939482126 |
| IDH1 | O75874 | Homo sapiens OX=9606 | 9.87676816158489 | 0.184527739086192 | -0.358249535050817 | 0.99980939482126 |

**One column need to be in common**

RNA-seq

| idx | diff_med_24h.logFC | diff_med_24h.AveExpr | diff_med_24h.t | diff_med_24h.P.Value | diff_med_24h.adj.P.Va |
|---|---|---|---|---|---|
| AAAS | 0.028183162940322 | 5.72297816146273 | 0.100490173895532 | 0.921125366636798 | 0.9917695300499 |
| AACS | 0.534149879708331 | 4.45269791483771 | 1.64919191277447 | 0.117377841874363 | 0.4509357956044 |
| AARS | 0.199342676608586 | 4.98757470941001 | 0.481363034947611 | 0.636367796688444 | 0.9279204702562 |
| AASDHPPT | -0.247161129015488 | 4.83935965937663 | -0.695725656217854 | 0.495959151345674 | 0.8313211270439 |
| ABCC1 | -1.23553413477472 | 2.70011516424125 | -1.65099903822641 | 0.117004717006166 | 0.4501955251607 |
| ABCD1 | 0.152322007640473 | 0.857479682917489 | 0.168071491533119 | 0.86850117891888 | 0.9917695300499 |
| ABCD3 | -0.281670225429062 | 6.41489681550556 | -1.32645373367621 | 0.202153780378643 | 0.5642399595980 |
| ABCE1 | 0.623063181229627 | 5.35006411905552 | 2.15407647213151 | 0.045802950961147 | 0.296402241764 |
| ABCF1 | 0.015813080864439 | 4.88613299788463 | 0.047478118279708 | 0.962682923075981 | 0.9917695300499 |

#### Supplementary materials S2: JSON settings objects used for generating figures in this study

In the GitHub release found at the DOI https://doi.org/10.5281/zenodo.4110102, the JSON objects with the settings used in OmicLoupe to generate the different visualizations presented in this study is present in the file “jsons_collection.txt”.

#### Supplementary materials S3: Case 1, dendrogram illustration of impact from software processing methods

Illustration of dendrogram clustering, highlighting clustering based on spike-in levels (upper) and the software used for processing (lower, pv: PeakView, os: OpenSWATH, sl: Skyline, du: DIA Umpire). Overall, methods cluster stronger than spike-in levels, with the exception of PeakView (pv) and OpenSwath (os).

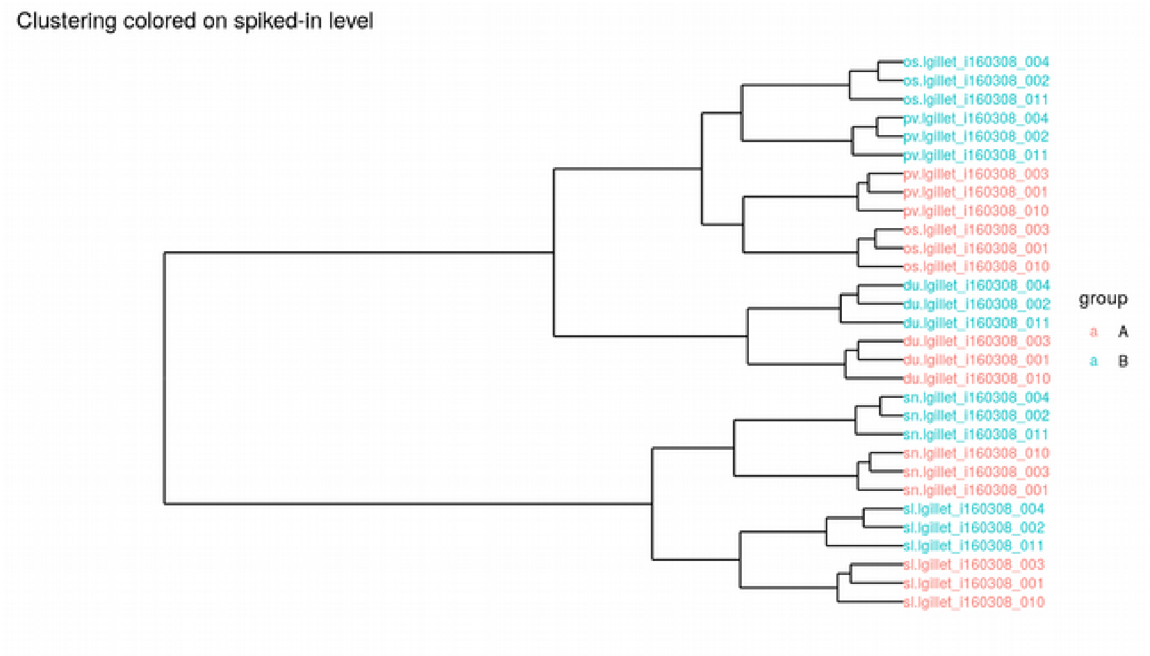


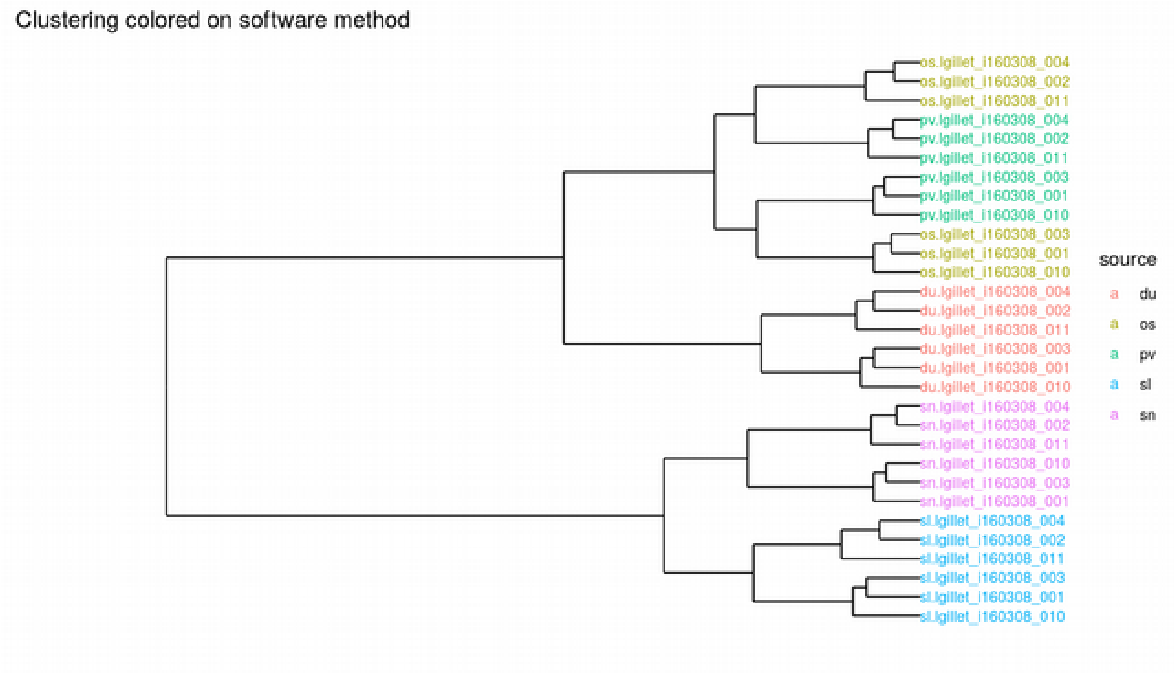

### Supplementary materials S4: Case 1, full statistical panel illustration of DIA Umpire compared to PeakView

Full grid statistical visualizations as obtained by OmicLoupe. Blue illustrates proteins which are differentially expressed (FDR < 0.05, log2 fold > 1) in both conditions and with the same fold direction, while green are diffentially expressed in both with reversed fold direction. Red are proteins only diffentially expressed in DIA Umpire, and yellow in PeakView.

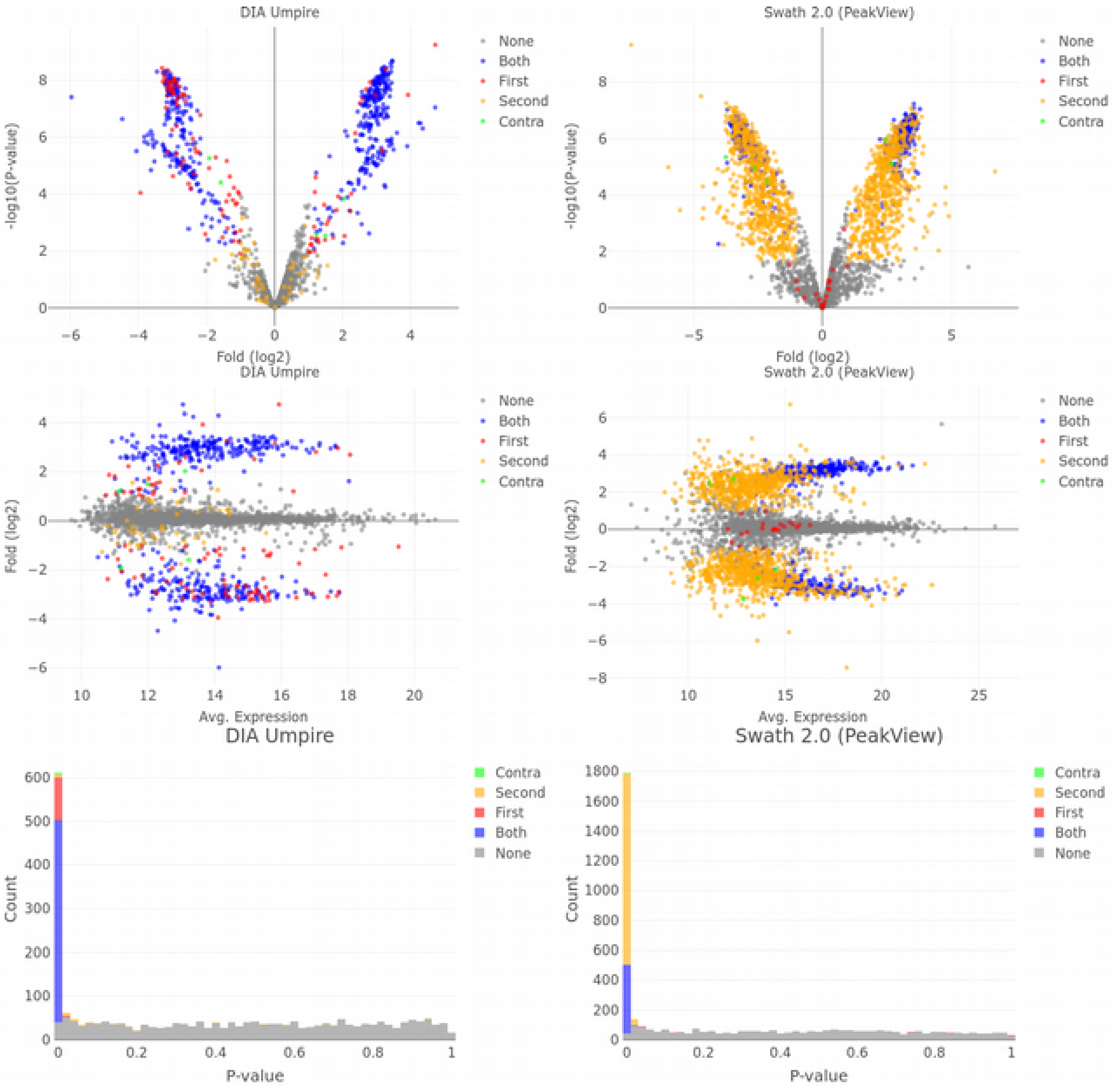

#### Supplementary materials S5: Case 2, PCA illustrations for subsets excluding non-cancerous data

Principal component analysis clustering of the multiomics dataset used in case 2. This illustrates how OmicLoupe can omit sub-groups of the data to better understand the clustering of the remaining samples. Here, the outlier group containing non-cancerous samples is omitted in both the proteomics and transcriptomics data.

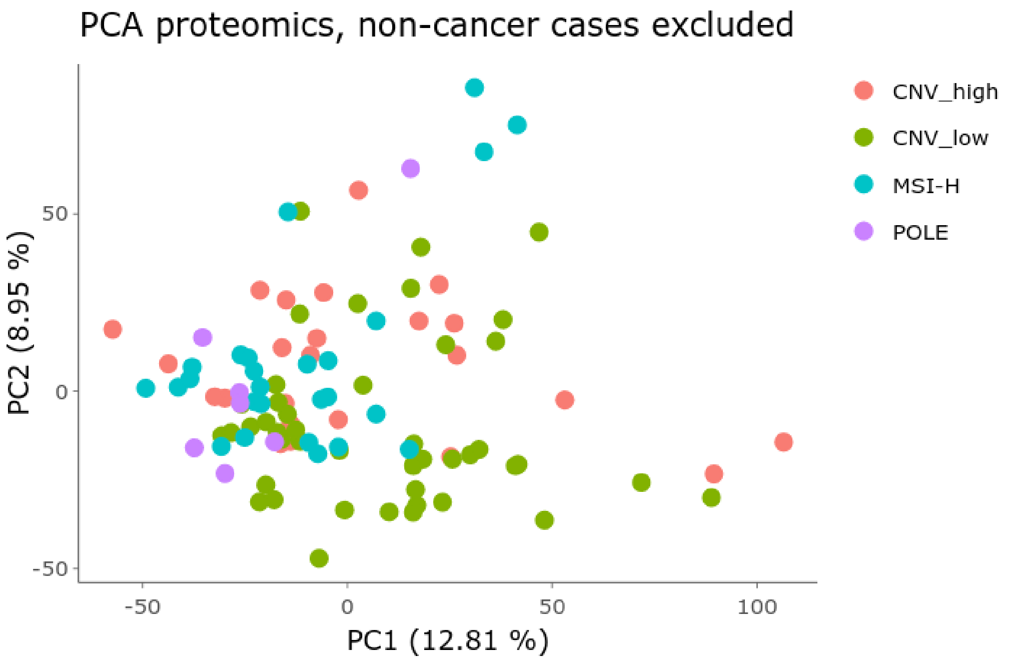


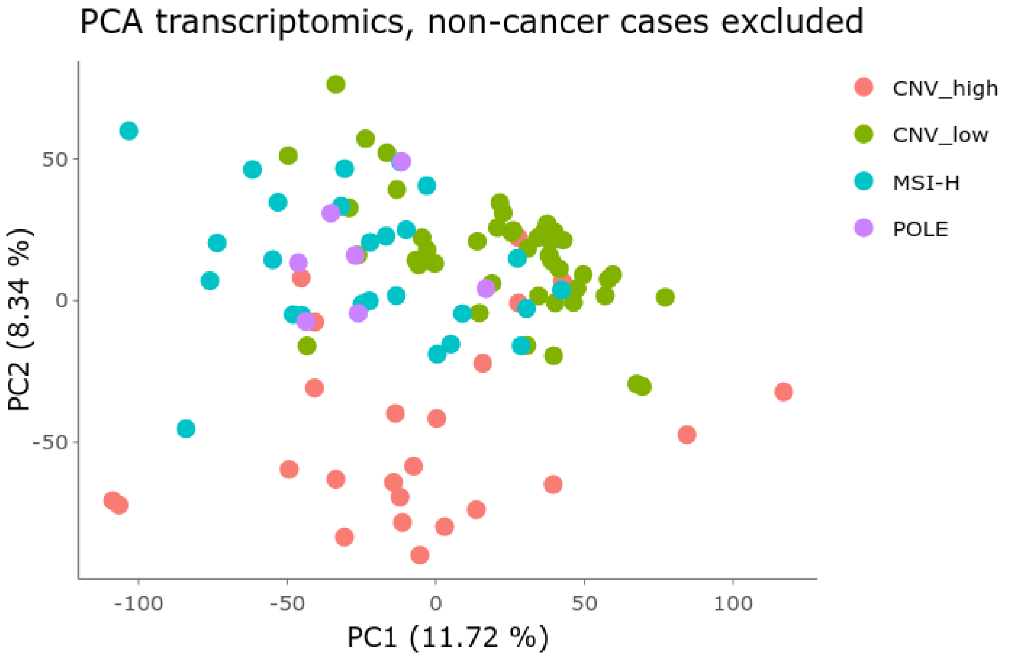

#### Supplementary materials S6: Case 3, proteomics PCA illustration of control- and infected samples

Principal component analysis inspection of control- and infected samples separately for the case 3 proteomics dataset. This highlights a reverse trend between control groups 6 hours and 24 hours after infection. Further, for infected samples belonging to the 24 hours after infection group is a strong outgroup, with a similar but weaker trend 10 hours after infection.

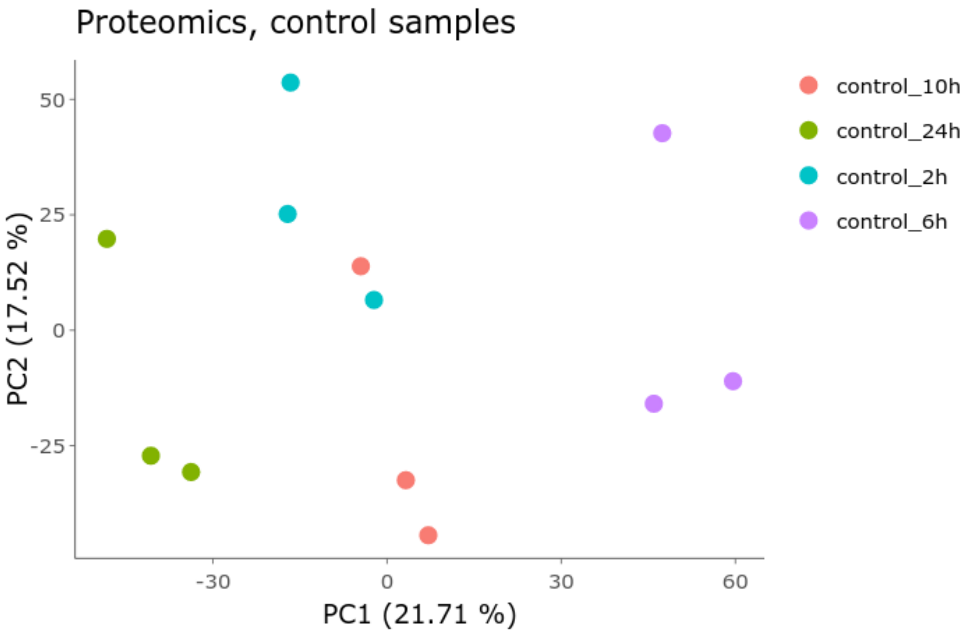


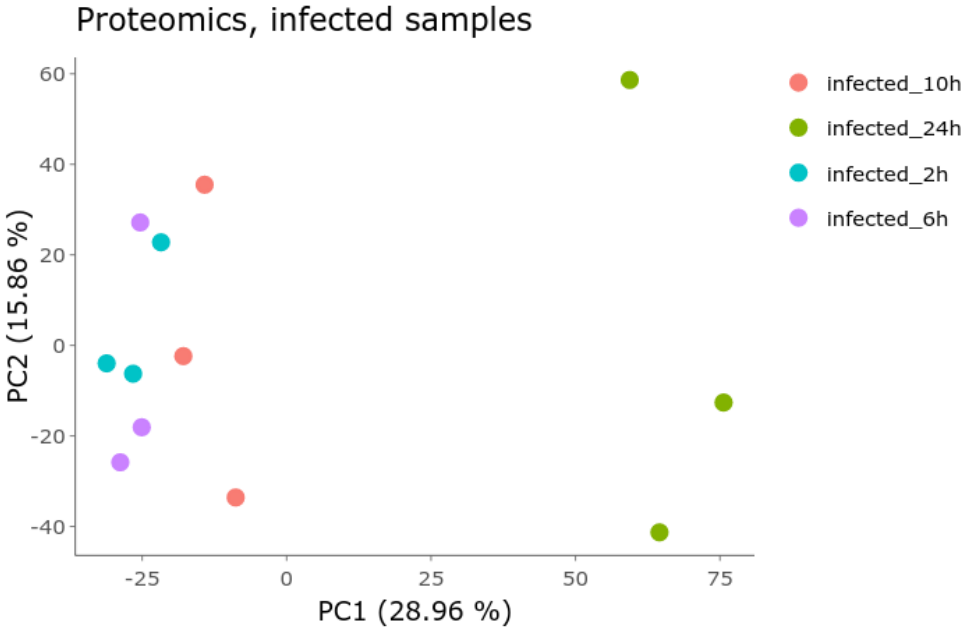

#### Supplementary materials S7: Case 3, PCA illustration of transcriptomics samples

Illustration of clustering in transcriptomics samples for case 3, showing the four first dimensions of the principal component analysis. Two outliers stands out - one SARS-CoV-2 sample 24 hours after infection on expansion medium (seen along PC2), and one SARS-CoV-2 after growing 60 hours on differentiation medium (seen along PC3).

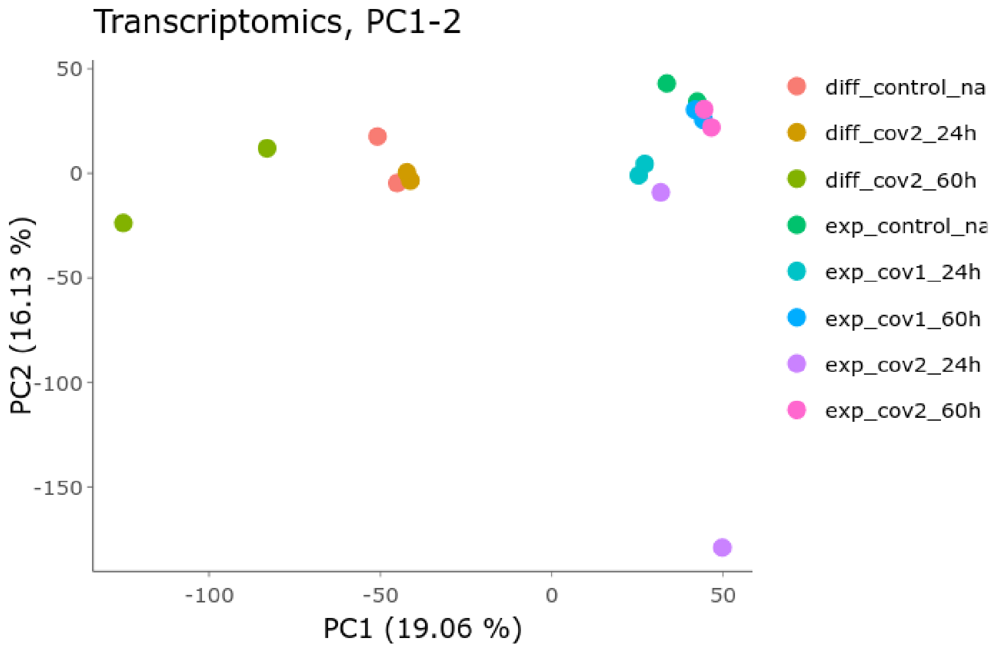


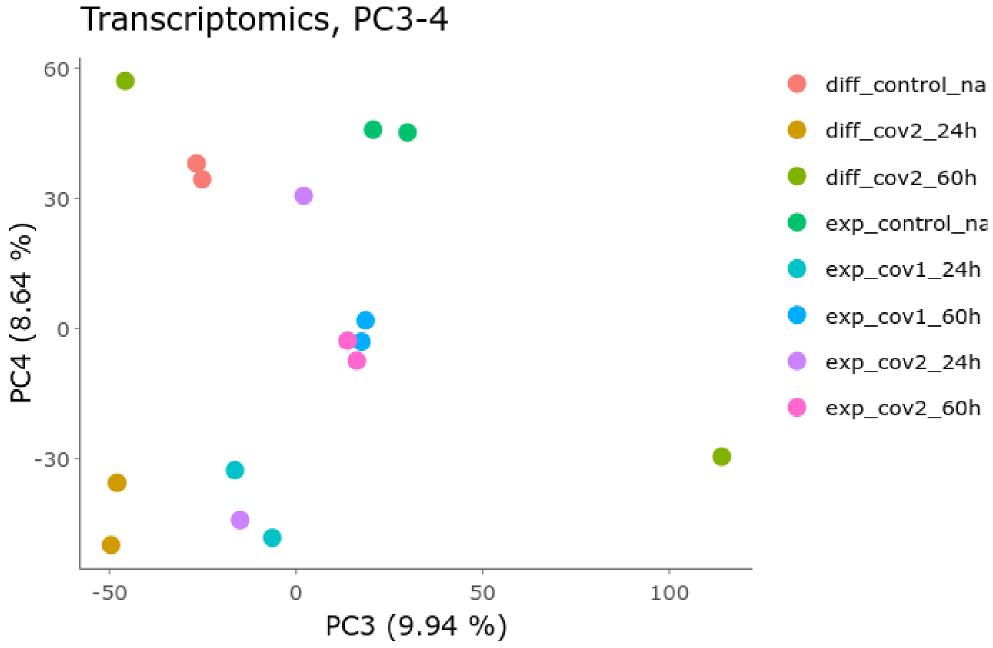

#### Supplementary materials S8: Analysis code for preprocessing datasets for study in OmicLoupe

In the GitHub release found at the DOI https://doi.org/10.5281/zenodo.4110102, the R code used for preprocessing the datasets into a format suited for OmicLoupe is present in the file "final_analysis.Rmd". Furthermore, the code is presented together with the output as obtained while running it in "final_analysis.html".
